# Fast-acting antidepressants trigger presynaptic BDNF release and structural plasticity at mouse mossy fiber-CA3 synapses

**DOI:** 10.64898/2026.01.02.697407

**Authors:** Irem L. Atasoy-Rodriguez, Kenneth W. Johnson, Kishan Patel, Haroon Arain, Syed Zaidi, Karl F. Herold, Teresa A. Milner, Hugh C. Hemmings, Jimcy Platholi

**Affiliations:** Department of Anesthesiology, Weill Cornell Medicine, New York, NY, USA; Department of Pharmacology, Weill Cornell Medicine, New York, NY, USA; Feil Family Brain and Mind Research Institute, Weill Cornell Medicine, New York, NY, USA

**Keywords:** fast-acting antidepressants, brain-derived neurotrophic factor, presynaptic NMDA receptors, mossy fiber terminals, CA3 dendritic spines

## Abstract

Major depressive disorder is associated with deficits in hippocampal synaptic plasticity that depend on brain-derived neurotrophic factor (BDNF) release from both axonal and dendritic compartments. Antidepressant efficacy requires enhanced BDNF signaling, thought to be mediated by drug-induced BDNF release from postsynaptic dendritic spines. Here, we show that fast-acting antidepressants rapidly trigger BDNF secretion from presynaptic terminals in hippocampal area CA3. At antidepressant-relevant concentrations, ketamine and its metabolite (2R,6R)-hydroxynorketamine (HNK) induced BDNF release within minutes from mossy fiber terminals of dentate granule neurons in rat hippocampal cultures, with no detectable secretion from dendritic spines. This antidepressant-evoked BDNF release required presynaptic NMDA receptors (preNMDARs). Conditional genetic deletion of preNMDARs from granule neurons abolished ketamine- and HNK-induced BDNF exocytosis in acute mouse hippocampal slices, establishing a presynaptic receptor mechanism for antidepressant-induced neurotrophin release. In CA3 pyramidal neurons that receive mossy fiber input, both compounds induced rapid remodeling of dendritic spines, resulting in increased spine density. Together, these findings identify presynaptic terminals as a previously unrecognized source of antidepressant-evoked BDNF release and establish a new cellular mechanism for the rapid synaptic effects of fast-acting antidepressants.

## Introduction

Ketamine, a widely used general anesthetic, has gained attention for its rapid and sustained antidepressant effects at low, subanesthetic doses (*1*). Clinical improvement can occur within one hour of a single infusion in individuals with major depressive disorder, including treatment-resistant cases (*2, 3*). These exceptionally fast actions are associated with activity-dependent synaptic plasticity (*4, 5*), yet the cellular sources and circuit mechanisms that initiate them remain incompletely understood.

Brain-derived neurotrophic factor (BDNF) is a key mediator of antidepressant-induced plasticity (*6–8*),. Transient increases in BDNF occur within ∼30 minutes across multiple brain regions (*5, 9–11*) and precede behavioral improvement, whereas sustained elevations are not required for long-term efficacy (*5*), suggesting that rapid, localized release may be critical for initiating antidepressant responses. Current understanding of ketamine-induced BDNF release has centered on postsynaptic dendritic compartments(*5, 12–15*). However, BDNF is also released from axonal terminals (*16–19*), where it can signal both locally and across synapses to regulate circuit plasticity (*20, 21*), suggesting that presynaptic sources contribute to its antidepressant actions.

Ketamine’s antagonism of NMDA receptors (NMDARs) has been proposed to facilitate activity-dependent BDNF release (*4, 5, 9*). Yet the specific NMDAR populations that mediate this effect remain unclear. Presynaptic NMDARs (preNMDARs) are well positioned to regulate neurotransmitter and neurotrophin release, and in the hippocampus, they facilitate BDNF secretion from mossy fiber terminals (*22*), large boutons of dentate gyrus (DG) granule neurons that project to CA3 pyramidal neurons and interneurons and play a central role in synaptic plasticity (*23*). In these hippocampal granule cell projections, BDNF is concentrated at exceptionally high levels (*23*) and is localized almost exclusively to presynaptic dense-core vesicles (*18*), consistent with activity-dependent release from axon terminals. Although this circuit has been implicated in the actions of several antidepressants (*24*), its role in ketamine-induced plasticity has not been directly examined, and whether presynaptic BDNF release contributes to rapid antidepressant mechanisms remains unknown.

Structural remodeling of dendritic spines is a hallmark of antidepressant action (*14, 25, 26*). Ketamine increases spine density in cortical and hippocampal neurons (*27, 28*), with structural changes persisting for weeks *in vivo* (*29*). Spinogenesis is required for the maintenance of antidepressant responses, although not for their initial emergence (*27*), suggesting that structural plasticity stabilizes circuit changes triggered by rapid signaling events. BDNF signaling regulates the cytoskeletal processes underlying spine formation (*30–32*), and given the high presynaptic stores of BDNF in mossy fiber terminals, acute release from these terminals could provide a mechanism for rapid, circuit-specific increases in dendritic spine density.

Here, using complementary rodent models, we show that fast-acting antidepressants rapidly evoke BDNF release from presynaptic mossy fiber terminals via preNMDAR signaling. Pharmacological manipulations and conditional genetic deletion demonstrate that ketamine-induced BDNF release occurs independently of NMDAR-mediated ion flux and postsynaptic NMDARs, whereas its metabolite hydroxynorketamine (HNK) requires both presynaptic and postsynaptic NMDARs. Consistent with a circuit-level consequence of this presynaptic signaling, ketamine and HNK produce rapid increases in dendritic spine density in CA3 pyramidal neurons. Together, these findings identify a previously unrecognized presynaptic source of antidepressant-evoked BDNF release and reveal a hippocampal synaptic mechanism that may initiate the rapid plasticity underlying the actions of fast-acting antidepressants.

## Results

### Ketamine and hydroxynorketamine (HNK) rapidly induce BDNF release from mossy fiber terminals

Both ketamine and its metabolite HNK elevate BDNF signaling to produce antidepressant effects (*1, 8*). To determine whether these fast-acting antidepressants directly trigger BDNF secretion from specific subcellular compartments (*17*), we measured BDNF release from presynaptic boutons and dendritic spines in dissociated rat hippocampal neurons expressing a pH-sensitive BDNF reporter fused to the C-terminus of BDNF (BDNF-SEP; Fig. 1). BDNF-SEP has been validated as properly processed, releasable, and biologically active (*17*). Axonal and dendritic pools were confirmed by NH_4_Cl alkalization at the end of each experiment (Fig. 1A). Acute application of ketamine (1 μM) and HNK (10 nM) evoked rapid BDNF release from presynaptic boutons within minutes and sustained asynchronous secretion throughout the 1 h confocal imaging period (Fig. 1, B to D and Supplementary Video 1). Based on bouton size and morphology, these structures were identified as mossy fiber terminals and further confirmed by immunocytochemistry (Fig. 1A and Supplementary Fig. 1). In contrast, neither ketamine nor HNK induced detectable BDNF release from dendritic spines within the same time window (Supplementary Fig. 2). Given that BDNF in the DG is required for antidepressant efficacy (*6, 18*), and that the mossy fiber pathway projecting to CA3 (Fig. 1E) contains the highest BDNF concentration in the CNS (*23*), these results identify mossy fiber terminals as a presynaptic source of antidepressant-evoked BDNF release engaged by fast-acting antidepressants.

**Fig. 1.**
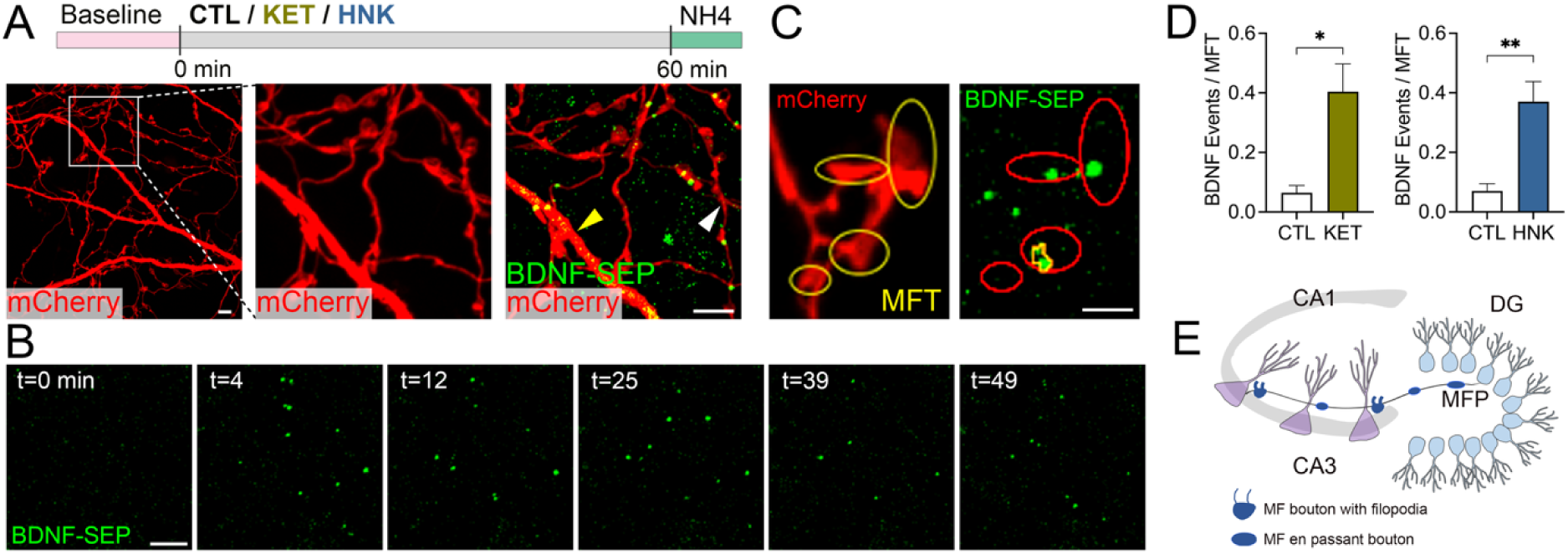
Rapid ketamine- and HNK-evoked release of BDNF from hippocampal mossy fiber terminals. (**A**) Rat hippocampal neurons (21 DIV) transfected with BDNF-SEP and mCherry were perfused with 1 μM ketamine (KET), 10 nM HNK, or control buffer (CTL) and imaged as indicated (*top*). Representative mCherry image of a control neuron showing dendritic arbors and axons (*left*). Inset: dendritic segment and single boutons before (*middle*) and after (*right*) NH_4_Cl alkalinization, revealing BDNF-SEP pools in axons (white arrowhead) and dendrites (yellow arrowhead). (**B**) Representative images of single BDNF exocytotic events (green dots) before (t = 0) and after (t = 4–49) ketamine application with individual BDNF release events selectively arising from mossy fiber terminals (MFTs) chosen for analysis (**C**, circles). (**D**) Quantification of BDNF exocytosis showing increased release following ketamine or HNK treatment (\**P* < 0.05 and \*\**P* < 0.01; Student unpaired *t*-test). (**E**) Schematic of the hippocampal mossy fiber pathway (MFP) illustrating dentate gyrus (DG) granule neurons with their *en passant* boutons and mossy fiber boutons synapsing onto CA3 pyramidal neurons. Data represent mean ± SEM, *n* = 5–10 cells and 302–825 MFTs selected for each condition. Scale bars: 5 μm (**A**, **B**) and 2 μm (**C**).

### NMDAR antagonism and presynaptic GRIN1 deletion reduce mossy fiber BDNF release induced by ketamine and HNK

Regulation of ligand- and voltage-gated channels (*1, 4*), including both direct and indirect modulation of NMDARs (*5, 33, 34*), has been proposed to converge on increases in BDNF signaling that drive synaptic plasticity required for antidepressant efficacy. At mossy fiber synapses, preNMDARs facilitate BDNF release underlying synapse-specific forms of plasticity (*22*), likely supported by their distinct subunit compositions (*35, 36*) that enable both ionotropic and metabotropic signaling (*37–39*). To verify NMDAR localization at mossy fiber terminals, we performed pre-embedding immuno-electron microscopy in mouse dorsal hippocampus (*40*) using a GluN3A-selective antibody (*41, 42*) (Fig 2, A to C). Silver-intensified immunogold (SIG) particles for GluN3A were significantly enriched at presynaptic mossy fiber terminals relative to postsynaptic CA3 dendritic spines (Fig. 2D), confirming robust presynaptic expression. We next tested whether NMDAR signaling contributes to rapid BDNF release induced by ketamine. Hippocampal neurons were preincubated (3 min) with either the noncompetitive NMDAR antagonist MK-801 (10 μM) or the competitive antagonist APV (50 μM) before ketamine application. APV abolished ketamine-evoked BDNF release (Fig. 2E), indicating that glutamate binding to NMDARs is required for this effect. In contrast, MK-801, an open channel blocker, failed to prevent ketamine-evoked BDNF release, even at saturating concentrations, and MK-801 alone did not induce detectable release (Fig. 2F). These findings indicate that ketamine-driven BDNF release depends on glutamate binding to NMDARs but not channel ion flux, consistent with a metabotropic NMDAR signaling.

**Fig. 2.**
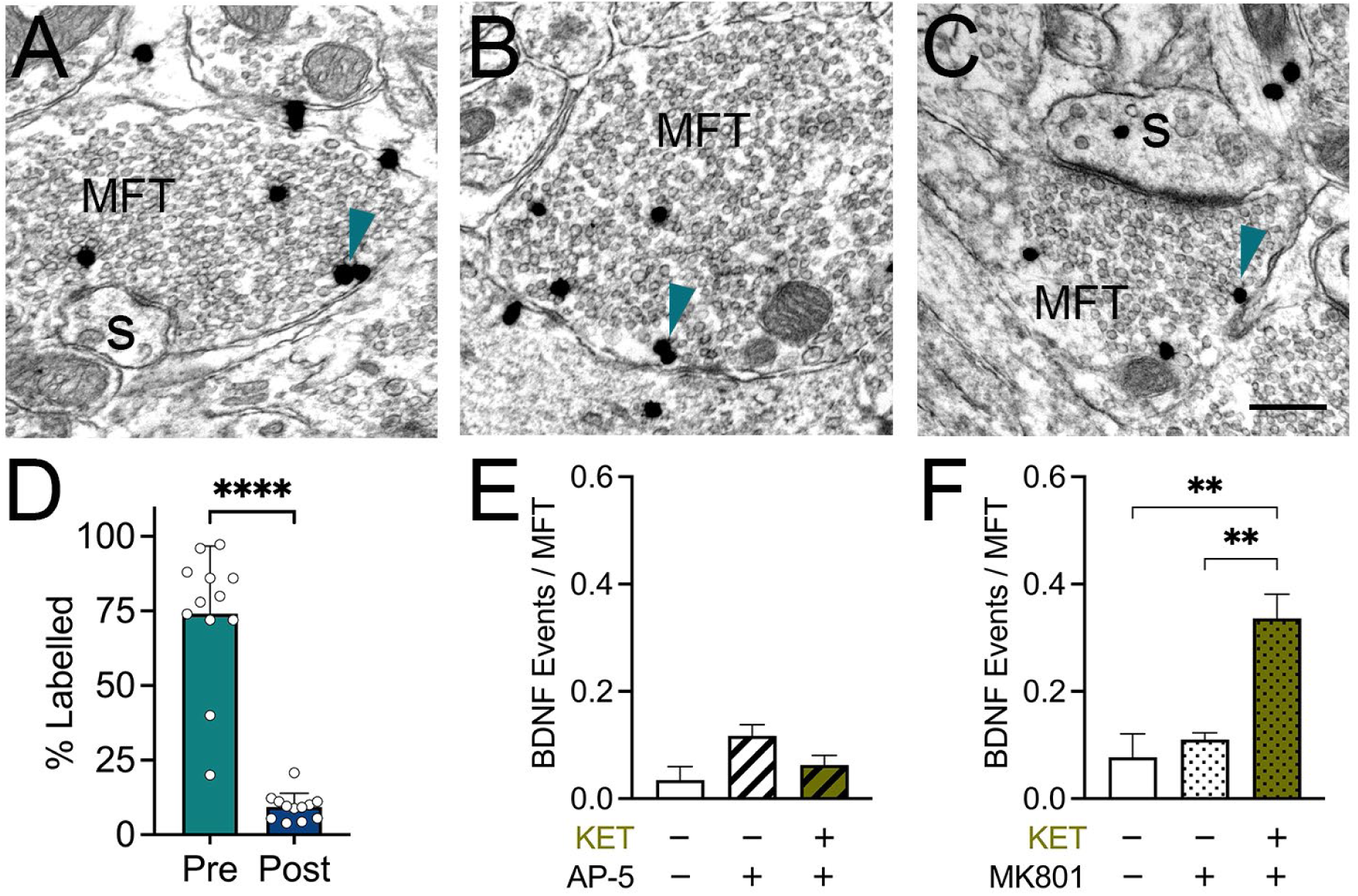
Anatomical and functional evidence for NMDAR regulation by ketamine and HNK. (**A–C**) Electron micrographs showing GluN3A SIG labeling (black dots) of mossy fiber terminals (MFTs; **A, B**) and dendritic spines (S; **C**). SIG particles were observed predominantly on or near (arrows) the plasma membrane in MFTs. (**D**) Quantification of GluN3A SIG particles demonstrating greater GluN3A expression in MFTs (Pre) compared to dendritic spines (Post). Data for 50 GluN3A-labeled MFTs from randomly selected micrographs were obtained from 12 mice (6 males and 6 females), with each data point representing one animal; *n* = 432 presynaptic profiles; *n* = 124 postsynaptic profiles. Data expressed as mean % labelled ± SEM (\*\*\*\**P* < 0.0001; Student unpaired *t*-test). Scale bar, 300 nm. (**E, F**) Quantification of BDNF exocytosis from MFTs showing that ketamine-evoked release is inhibited by preincubation with APV (50 μM) (**E**) but not MK-801 (10 μM) (**F**). \*\**P* < 0.01; one-way ANOVA. Data represent mean ± SEM, *n* = 3–7 cells and 222–596 MFTs analyzed per condition.

Because bath-applied antagonists affect both axonal and dendritic NMDARs, we selectively deleted the obligatory NMDAR subunit *GRIN*1 from DG granule neurons in *Grin1*^flox^ mice to assess the specific contribution of preNMDARs. Cre-dependent mice (postnatal days 23–35) received unilateral DG injections of AAV5-CaMKII-mCherry-cre and AAV-DJ-DIO-BDNF-pH to selectively delete *Grin1* and express the BDNF-pH reporter, respectively (Fig. 3A). Wild-type mice receiving the same viral injections served as controls. Robust expression of pHluorin (GFP) and mCherry in DG granule neurons and mossy fiber terminals was confirmed in acute hippocampal slices four weeks after injection (Fig. 3, B to D), and only slices exhibiting strong expression of both reporters were analyzed.

**Fig. 3.**
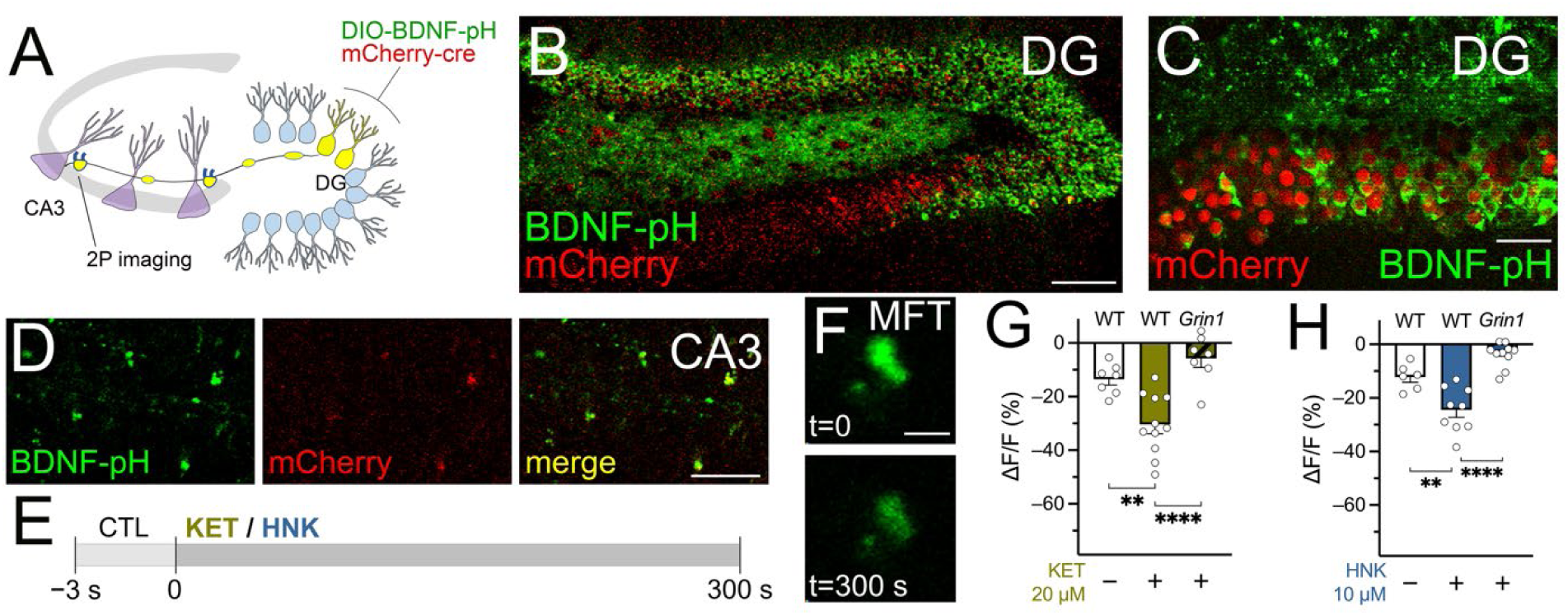
preNMDARs are required for ketamine- and HNK-evoked BDNF release from mossy fiber terminals. (**A**) Schematic illustrating viral transduction of DG granule neurons with mCherry (red) and BDNF-pH (green), resulting in co-labeling of MFTs at imaging sites in CA3 (yellow). (**B–D**) Representative images from acute hippocampal slices from *Grin1*^flox^ mice showing robust mCherry-cre expression (red; deleting GluN1 and enabling Cre-dependent BDNF-pH expression) in DG (**B, C**) and merged BDNF-pH + mCherry labeling in MFT in CA3 (**D**). (**E**) Experimental timeline showing baseline recordings in ACSF (CTL) followed by perfusion with 20 μM ketamine (KET) or 10 μM HNK. (**F**) Representative images of BDNF-pH fluorescence in MFTs at baseline (t = 0) and after 300 s of ketamine exposure, showing decreased fluorescence corresponding to BDNF release. (**G, H**) Quantification of BDNF release from MFTs demonstrating that ketamine-evoked (**G**) and HNK-evoked (**h**) BDNF secretion is abolished in slices lacking NMDAR function in DG granule neurons. (\*\**P* < 0.01, \*\*\*\**P* < 0.0001; one-way ANOVA with Tukey *post hoc* test). Data are mean ± SEM, *n* = 6–7 animals and 7–11 slices per condition. Scale bars:100 μm (**B**), 30 μm (**C, D**), and 5 μM (**F**).

Using two-photon imaging, we found that ketamine (20 μM) and HNK (10 μM), concentrations previously shown modulate hippocampal plasticity in cultured neurons and slices (*43, 44*), produced a marked reduction in BDNF release from mossy fiber terminals in acute slices from *Grin1*-deleted mice within 5 minutes of treatment (Fig. 3, E to H). Together, these findings indicate that NMDAR signaling, particularly in DG-derived mossy fiber terminals, is required for rapid BDNF release induced by ketamine and HNK.

### Postsynaptic CA3 NMDARs selectively mediate HNK-evoked BDNF release

To determine whether postsynaptic NMDARs participate in ketamine- and HNK-induced BDNF release, we conditionally deleted *Grin1 from* CA3 pyramidal neurons in *Grin1*^flox^ mice. Cre-dependent mice (postnatal days 23-35) received unilateral injections of AAV5-CaMKII-mCherry-cre into area CA3 to restrict *Grin1* deletion to neurons receiving mossy fiber input. Because CA3 pyramidal cell bodies lie in close proximity to DG axons, complete avoidance of retrograde viral spread is not technically feasible, raising the possibility of partial presynaptic knockdown. To distinguish presynaptic from postsynaptic contributions, we simultaneously injected a non-Cre-dependent AAV-DJ-BDNF-pH into CA3 to express the BDNF-pH reporter independently of *Grin1* status (Fig. 4A). This strategy yielded three fluorescent readouts: *Grin1* deletion (mCherry; red), BDNF-pH expression (pHluorin; green), and co-expression of BDNF-pH with *Grin1* deletion (mCherry + pHluorin; yellow) (Fig. 4B). To ensure that fluorescence measurements reflected release from mossy fiber terminals synapsing onto CA3 neurons lacking postsynaptic NMDARs, analyses were restricted to slices exhibiting strong mCherry expression in CA3 somata but pHluorin-only expression in mossy fiber terminals (Fig 4C). Under these conditions, ketamine (20 μM) evoked presynaptic BDNF release, indicating that somatodendritic NMDARs in CA3 are not required for acute ketamine actions. In contrast, HNK (10 μM) evoked-BDNF release was significantly reduced in slices lacking postsynaptic NMDARs (Fig. 4, D and E). These findings indicate that HNK, but not ketamine, additionally requires postsynaptic NMDAR signaling to drive rapid BDNF release at mossy fiber-CA3 synapses.

**Fig. 4.**
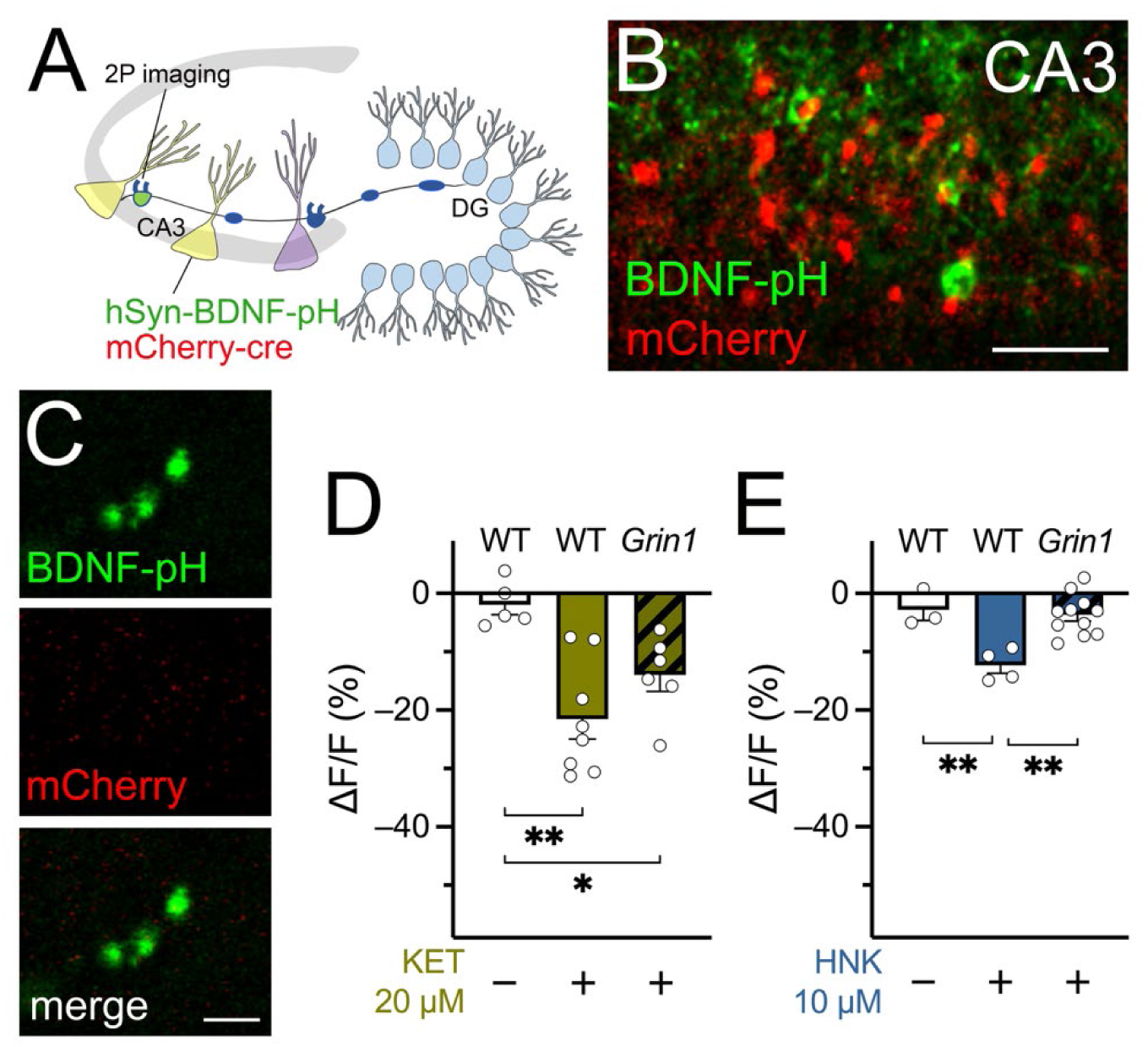
Postsynaptic NMDARs are required for HNK-evoked, but not ketamine-evoked, BDNF release from mossy fiber terminals. (**A**) Schematic illustrating viral transduction of CA3 pyramidal neurons with mCherry (red) and BDNF-pH (green), resulting in non-overlapping BDNF-pH-labeled MFT in CA3 (green). (**B, C**) Representative images from acute hippocampal slices from *Grin1*^flox^ mice showing strong mCherry-cre expression (red; deleting GluN1 in CA3 neurons) and non-Cre-dependent BDNF-pH expression (green) in CA3 (**B**), with BDNF-pH-labeled MFTs visualized in CA3 (**C**). (**D, E**) Quantification of BDNF release from MFTs demonstrating that ketamine-evoked secretion (**D**) persists in slices lacking NMDAR function in CA3 neurons, whereas HNK-evoked release (**E**) is abolished. (\**P* < 0.05, \**P* < 0.01, \*\**P* < 0.01; one-way ANOVA with Tukey *post hoc* test). Data are mean ± SEM, *n* = 5–8 animals and 3–11 slices per condition. Scale bars: 30 μm (**B**) and 5 μm (**C**).

### Ketamine and HNK rapidly increase dendritic spine density in CA3 pyramidal neurons

Structural plasticity of dendritic spines is a hallmark of antidepressant action (*26, 27*), yet the initial drug-evoked changes within hippocampal CA3 remain poorly defined. We therefore examined whether rapid alterations in CA3 apical dendritic spines, which receive synaptic input from mossy fiber terminals, coincide temporally with the acute burst of BDNF release triggered by ketamine or HNK. Using two-photon microscopy, we imaged eGFP-labeled CA3 apical dendrites in acute slices from Thy1-eGFP mice before and after 30 min of drug exposure (Fig. 5). Based on regional eGFP expression, analyses were largely restricted to second-order distal dendrites extending from the primary apical branch (Fig. 5a). Ketamine (20 µM) or HNK (10 µM) each produced rapid increases in CA3 dendritic spine density (Fig. 5, b to f). At the level of individual spines, both gains and losses were observed across dendrites, indicating dynamic remodeling; however, spine formation predominated, resulting in a net increase in overall spine density. These findings demonstrate that fast-acting antidepressants rapidly enhance structural connectivity in CA3 pyramidal neurons.

**Fig. 5.**
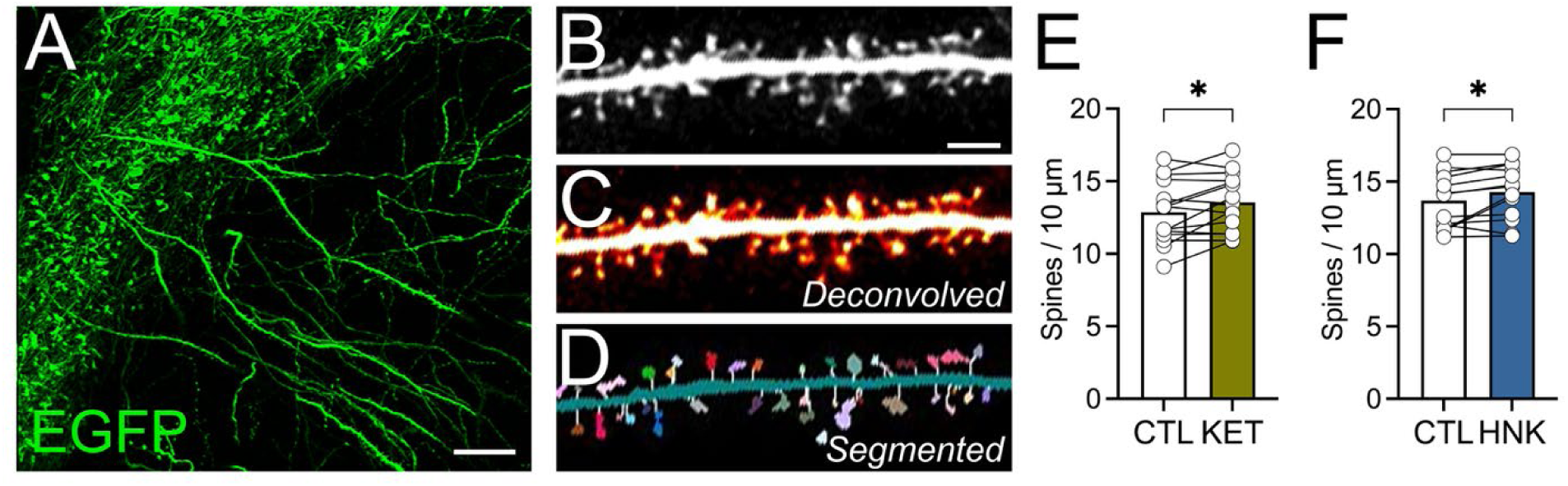
Ketamine and HNK rapidly remodel CA3 dendritic spines increasing overall spine density. (**A**) Representative image of hippocampal slice from Thy1-eGFP showing dendritic arbors and spines. (**B–D**) Image analysis pipeline illustrating a dendritic segment before processing (**B**), after deconvolution (**C**), and following automated segmentation (**D**). (**E, F**) Quantification of CA3 spine dynamics showing that 30 min treatment with ketamine (KET; **E**) or HNK (**F**) significantly increased spine density (\**P* < 0.05; Student paired two-tailed *t*-test). *n* = 5–8 animals, 14–15 slices, and 826–926 spines per condition. Scale bars = 30 μm (**A**) and 5 μm (**B–D**).

## Discussion

BDNF-dependent synaptic plasticity is essential for the onset of antidepressant actions, yet direct release of BDNF by antidepressant compounds has not been demonstrated. Here we show that ketamine and its metabolite HNK trigger rapid, selective BDNF release from presynaptic mossy fiber terminals within minutes of application. These terminals are enriched in presynaptic NMDARs, which we identify as necessary for BDNF release by both drugs, whereas postsynaptic NMDARs on CA3 neurons are specifically required for HNK-, but not ketamine-induced, release. In parallel, ketamine and HNK each drive rapid structural plasticity in CA3 dendritic spines, resulting in a net increase in spine density. This enhancement of hippocampal spine structure, coupled with drug-evoked BDNF release from mossy fiber terminals, reveals a previously unrecognized synaptic mechanism engaged at the onset of fast-acting antidepressant action.

Numerous studies have linked antidepressant efficacy to increases in hippocampal BDNF levels with subregion specificity. Selective loss of BDNF in the DG, but not CA1, abolishes behavioral responses to classical antidepressants, and deletion of its receptor, tropomyosin receptor kinase B (TrkB), in the DG blocks antidepressant-induced neurogenesis (*45*). For fast-acting agents such as ketamine, a single infusion into the medial prefrontal cortex or hippocampal DG and CA3 is sufficient to elicit antidepressant-like effects (*46, 47*). Within the DG, specific cellular substrates have been identified: activity of immature granule neurons is required for the rapid behavioral response to ketamine, whereas adult neurogenesis supports its sustained effects (*48, 49*). Although the DG-dependent role of BDNF in ketamine action has been well established, direct evidence of BDNF release triggered by ketamine or its metabolite HNK has been lacking. Here we show that both compounds acutely evoke BDNF exocytosis from mossy fiber terminals of DG granule neurons, providing a synapse-specific mechanism that links DG activity to CA3 circuit plasticity. These findings suggest that early BDNF release at DG-CA3 synapses may represent a key initiating step in the neuropharmacological actions of ketamine and HNK.

Ketamine is a noncompetitive NMDA antagonist, and modulation of NMDAR activity regulates BDNF signaling through various cell-type-specific mechanisms. In the PFC, tonic inhibition of NMDARs on GABAergic interneurons leads to disinhibition of pyramidal neurons, increasing dendritic spine calcium and promoting BDNF release via mTOR signaling (*9, 50*). In the hippocampus, attenuation of resting NMDAR activity on CA1 pyramidal neurons by ketamine or HNK enhances local BDNF translation through eukaryotic elongation factor 2 (eEF2) kinase-dependent mechanisms, thereby increasing synaptic drive and homeostatic plasticity (*5, 51, 52*). More recently, both classical and fast-acting antidepressants have been shown to directly bind the BDNF receptor TrkB (*53*), facilitating BDNF signaling independently of NMDAR blockade. These findings indicate that multiple forms of plasticity (*50, 52, 54*) and cell-type-specific NMDAR mechanisms contribute to acute and sustained phases of antidepressant action. BDNF secretion can occur at both presynaptic and postsynaptic sites (*17, 18, 55*), and our results identify a synapse-specific presynaptic source engaged by ketamine and HNK. Presynaptic NMDARs at mossy fiber terminals were required for rapid BDNF release induced by both drugs, whereas postsynaptic NMDARs on CA3 dendrites were necessary only for HNK-evoked release. As mossy fiber terminals synapse onto both CA3 pyramidal neurons and interneurons, contributions from NMDARs on local interneurons cannot be excluded and remain to be examined. Mossy fiber-CA3 synapses support diverse forms of short-term (*56, 57*), long-term (*56, 58*), and homeostatic plasticity (*59, 60*), placing preNMDARs at mossy fiber terminals as key regulators of antidepressant-evoked BDNF secretion within the CA3 circuit.

BDNF exocytosis was confined to mossy fiber terminals and was not detected from dendritic spines within 1 h of ketamine or HNK treatment. This contrasts with prior studies reporting rapid increases in postsynaptic BDNF translation in hippocampal dendrites mediated by eEF2K-eEF2 signaling (*5, 61*). Similarly, we did not observe increased BDNF release with MK-801, an NMDA antagonist that produces acute antidepressant-like behavioral responses (*5, 62, 63*), consistent with earlier reports (*5, 64*). One potential explanation is that NMDARs containing GluN1/GluN3A subunits, which are enriched at mossy fiber terminals, are largely insensitive to open-channel blockers such as Mg^2+^, memantine, or MK-801 (*41, 65, 66*). More broadly, these differences may reflect variation in experimental preparation, brain subregion (*67*), or methods used to quantify BDNF. pHluorin-based reporters measure secretion events rather than total protein levels and may not directly correlate with expression or translation. Our finding that BDNF release originates from mossy fiber terminals is consistent with evidence that BDNF transcription and translation occur in granule neurons (*68*) and that activity promotes its accumulation in presynaptic terminals (*23, 68*). However, the mechanisms governing compartment-specific localization and secretion of BDNF remain poorly understood (*69, 70*). Notably, the requirement for postsynaptic NMDARs in HNK-evoked, but not ketamine-evoked, BDNF release suggests that postsynaptic receptor activation can retrogradely influence presynaptic secretion, potentially through rapid trans-synaptic signaling pathways (*71–73*). Together, these observations raise the possibility that fast-acting NMDAR antagonists differentially engage presynaptic *and* postsynaptic mechanisms depending on synapse type and temporal stage of drug action.

In mossy fiber terminals, evoked and spontaneous neurotransmitter release are differentially regulated by glutamate binding to preNMDARs. Activity-dependent release requires Ca^2+^ influx through the receptor, whereas spontaneous transmission can occur via metabotropic NMDAR signaling independent of ion flux, mediated by conformational changes that engage Rab3 interacting molecule 1 (RIM1) or c-Jun N-terminal kinase 2 (JNK2) pathways (*37*). At resting membrane potentials, the NMDAR pore is largely blocked by extracellular magnesium (Mg^2+^) (*74*); however, this block is incomplete under physiological conditions, permitting small residual currents that can be inhibited by ketamine or MK-801 (*75*) through mechanisms involving transient Mg^2+^ block (*51, 75*) or interactions at distinct binding sites (*75, 76*). Our observation that ketamine elicits BDNF release from mossy fiber terminals is consistent with reports that ketamine suppresses spontaneous NMDAR activity at rest (*51, 75*). Notably, when residual NMDAR currents were first blocked with MK-801, subsequent ketamine treatment still triggered robust BDNF release, whereas occluding glutamate binding with APV prior to ketamine attenuated this effect. These findings indicate that receptor activation by glutamate, but not ion flux through the channel, is required for ketamine-evoked BDNF exocytosis, consistent with a metabotropic mode of preNMDAR signaling that enhances rapid, synapse-specific BDNF release.

Both NMDARs and AMPA receptors (AMPARs) have been implicated in the antidepressant actions of ketamine (*1, 77*). For HNK, in addition to its reported inhibition of NMDARs (*51*), early and sustained activation of AMPARs independent of NMDAR blockade has been described (*1, 43*), suggesting distinct channel-gating mechanisms. In dissociated cultures, ketamine was applied at 1 μM, approximating clinically relevant free plasma concentrations(*71*), whereas HNK was applied at 10 nM, consistent with previous studies (*78*). In acute hippocampal slices, higher concentrations of ketamine (20 μM) and HNK (10 μM) were used to account for tissue binding and diffusion, in line with earlier slice studies examining rapid synaptic and structural plasticity (*43, 44, 51*); these concentrations were empirically verified to evoke robust BDNF release. Under these conditions, HNK-evoked BDNF release required both presynaptic and postsynaptic NMDARs signaling, consistent with a mechanism that engages multiple receptor populations within the mossy fiber-CA3 circuit.

NMDARs containing GluN3A subunits exhibit distinct biophysical, synaptic signaling, and subcellular localization properties compared with canonical receptors composed of GluN1 and GluN2(A-D) subunits (*41*). These receptors show reduced single-channel conductance, lower calcium permeability, and diminished sensitivity to Mg^2+^ block at hyperpolarized membrane potentials (*41, 79*), features that support tonic activity under resting conditions. Consistent with this profile, we confirm GluN3A expression in mossy fiber terminals and extend prior work demonstrating that preNMDARs facilitate BDNF release at mossy fiber synapses (*22*). The presence of GluN3A-containing NMDARs at mossy fiber terminals provides a plausible molecular basis for the sensitivity of presynaptic BDNF release to ketamine and HNK. In neurons, GluN3A can assemble with GluN2A or GluN2B subunits (*80, 81*), both implicated in ketamine’s cell type-specific actions in excitatory and inhibitory circuits (*13, 50, 62*). Although GluN2B-selective antagonists produce antidepressant-like effects in preclinical models (*13, 50*), these findings have not translated to clinical efficacy(*1*), underscoring the importance of defining subunit-specific mechanisms. Whether GluN3A-containing receptors at mossy fiber terminals preferentially assemble with GluN2A or GluN2B, and how such assemblies influence sensitivity to ketamine or HNK, remains unknown. To our knowledge, this study provides the first evidence implicating GluN3A-containing NMDARs at mossy fiber terminals as a molecular substrate for fast-acting antidepressant-evoked BDNF release. This observation is consistent with prior work linking GluN3A signaling to enhanced presynaptic release probability, long-term potentiation, and improved cognitive function (*82, 83*).

Overall spine density in CA3 increased after 30 minutes of ketamine or HNK treatment; however, individual dendritic branches exhibited heterogeneous changes in spine number, indicating that acute structural remodeling was spatially selective rather than uniform across the dendritic arbor (*27*). Such branch-specific remodeling suggests that fast-acting antidepressants refine local synaptic connectivity rather than producing indiscriminate growth. Consistent with this framework, hippocampal dendritic spines are functionally clustered (*84, 85*), and the likelihood that a spine is gained, stabilized, or eliminated may depend on local synaptic activity and molecular composition. Spines receiving strong glutamatergic or neurotrophic drive, including those enriched for BDNF-or glutamate-responsive signaling pathways, may be preferentially stabilized or expanded, whereas less active, labile, or silent spines may remain unchanged or be selectively pruned.

Future studies should define how ketamine and HNK differentially engage presynaptic and postsynaptic mechanisms to regulate mossy fiber-CA3 signaling and structural plasticity. Targeted manipulation of NMDARs and AMPARs in CA3 pyramidal neurons and local interneurons, using electrophysiology, glycine-site antagonists such as 7-chlorokynurenic acid (7-CKA), and conditional genetic approaches, will be necessary to directly assess receptor activation or inhibition and to distinguish metabotropic from ionotropic NMDAR signaling in shaping BDNF release. Parallel *in vivo* studies will be required to determine whether these synaptic mechanisms translate into antidepressant-like behavioral effects. At the molecular level, experiments in *Grin3a*-null mice (*83*) and the use of GluN3-selective antagonists (*86*) will clarify the specific contribution of GluN3A-containing NMDAR assemblies to ketamine- and HNK-evoked BDNF exocytosis. Finally, integrating analyses of spine subclass and dendritic branch-order, key determinants of synaptic strength, stability, and plasticity (*87, 88*), with targeted manipulation of BDNF and glutamate signaling (*88, 89*) will elucidate how activity at mossy fiber terminals drives spatially selective spine remodeling. Such approaches will be essential for linking subcellular structural plasticity to circuit function and the behavioral efficacy of fast-acting antidepressants.

## Materials and Methods

### Animals

Sprague Dawley rats were obtained from Charles River Laboratories (Wilmington, MA, USA). Homozygous mouse lines were acquired from Jackson Laboratory (Bar Harbor, ME; Tg(Thy1-EGFP)MJrs/J-#007788, B6.129S4-Grin1*^tm2Stl^*/J-#005246, and C57BL/6J-#000664) and bred in-house to maintain colonies. All animals were group-housed under a 12h light/dark cycle with free access to food and water. All animal procedures conformed to the National Institutes of Health *Guide for the Care and Use of Laboratory Animals* and were approved by the Weill Cornell Medicine Institutional Animal Care and Use Committee (IACUC). Animal handling and reporting adhered to ARRIVE guidelines.

### Viruses and cDNA constructs

AAV vectors were obtained from UNC Vector Core (University of North Carolina at Chapel Hill, Chapel Hill, NC), including AAV5-CaMKII-mCherry, AAV5-CaMKII-mCherry-cre, control AAV-CaMKIIa-GFP, and AAV-CaMKIIa-GFP-Cre. Additional viral vectors (AAV-hSyn-BDNF-pHluorin, cre-dependent AAV-DJ-DIO-BDNF-pHluorin) were custom-produced by the UNC vector core using plasmids generously provided by Dr. Hyungju Park (Korea Brain Research Institute, Daegu, Korea) from UNC Vector Core (AAV-hSyn-BDNF-pHluorin, cre-dependent AAV-DJ-DIO-BDNF-pHluorin). The BDNF-SEP plasmid was obtained from Addgene (#83955; Watertown, MA).

### Materials

Ketamine, hydroxynorketamine, and all other chemicals were obtained from Sigma-Aldrich (St. Louis, MO). Isoflurane was purchased from Henry Schein Medical (Melville, NY). All additional reagents were sourced as indicated.

### Antibodies

An affinity-purified polyclonal guinea pig anti-Zn_3_T antibody, previously characterized (*90*), was purchased from Synaptic Systems (Goettingen, Germany). A polyclonal rabbit anti-GluN3A antibody (recognizing amino acids 1098–1113 of rat GluN3A), previously validated in (*35*), was also used.

### GluN3A Immuno-electron microscopy

Electron microscopic immunolocalization of GluN3A was performed using the pre-embedding silver intensified immunogold method as described previously (*40*). Briefly, mice were deeply anesthetized with pentobarbital (150 mg/kg; i.p.) and perfused through the aorta sequentially with: 1) ∼5 ml 0.9% NaCl containing 1,000 units/ml heparin; and 2) 30 ml 3.75% acrolein and 2% paraformaldehyde in 0.1 M phosphate buffer (PB; pH 7.4). The brain was post-fixed in in 2% acrolein and 2% paraformaldehyde in PB for 30 min. and then the region containing the hippocampal formation (HF) was cut into a 5-mm-thick coronal block using a brain mold. Sections (40 μm thick) were cut through the HF on a vibrating microtome (VT1000 Leica Microsystems), and the sections were stored in 24-well plates in cryoprotectant solution (30% sucrose, 30% ethylene glycol in PB) at −20°C until use. Two dorsal hippocampal sections per mouse were selected from each experimental condition and coded by hole-punches in the cortex. The sections were then placed in a single container and processed together through all immunocytochemical procedures to ensure identical labeling conditions. Sections were incubated in 1% sodium borohydride in PB for 30 min to neutralize reactive aldehydes, then rinsed in PB ∼10 times until gaseous bubbling ceased.

Sections were rinsed in 0.1 M Tris saline (TS; pH 7.6) followed by an incubation in 0.5% BSA in TS for 30 min to reduce nonspecific labeling. Sections were then incubated in rabbit anti-GluN3A (1:50) in 0.1% BSA in TS for 1 day at RT and then 1 day at 4°C. Sections were rinsed in TS followed by washing buffer [0.1 M phosphate-buffered saline (PBS) with 2% gelatin and 0.1% BSA], and incubated overnight at 4°C in a 1:50 dilution of donkey anti-rabbit IgG conjugated to 1-nm gold particles [Electron Microscopy Sciences (EMS) Cat# 810.311] in 0.01% gelatin and 0.08% BSA. All primary and secondary antibody incubations were carried out at 145 rpm, and all rinses were conducted at 90 rpm on a rotator shaker. Sections were rinsed in PBS, post-fixed in 2% glutaraldehyde in PBS for 10 min, then rinsed in PBS and in 0.2 M sodium citrate buffer (pH 7.4). The conjugated gold particles were enhanced using silver solution (SEKL15 Silver enhancement kit, Prod No. 15718 Ted Pella Inc.) for 9 min. Sections were fixed in 2% osmium tetroxide for 1 h, dehydrated in increasing ethanol concentrations to propylene oxide, and embedded in EMBed 812 (EMS) between two sheets of Aclar plastic (Honeywell, Pottsville, PA). Ultrathin sections (70 nm) of the CA3 were cut with a Diatome diamond knife (EMS) on a Leica UCT6 ultratome and collected on 400 mesh thin-bar copper grids (T400-Cu, EMS). The grids were counterstained with Uranylless^TM^ (catalog #22409, EMS) and lead citrate (catalog #22410 EMS). Specimens were imaged on a Hitachi HT7800 transmission electron microscope.

### Hippocampal neuron culture and transfection

Primary rat hippocampal neurons were cultured as described previously (*91*). Briefly, hippocampi were dissected from embryonic day 16–18 (E16–18) Sprague Dawley rat embryos, dissociated using papain, and plated on Lab-Tek II chambered borosilicate coverglass (4.0 cm^2^/well; NUNC, Rochester, NY) at a density of 200 cells/mm^2^. Cells were maintained in Neurobasal medium (GIBCO, Grand Island, NY) supplemented with SM1 (Stem Cell Technologies, Vancouver, BC, Canada) and 0.5 mM L-glutamine (Sigma-Aldrich, St. Louis, MO). At 16 days *in vitro* (DIV), cultures were co-transfected using calcium phosphate precipitation with 6–10 µg of total DNA containing a CAG-driven mCherry plasmid and BDNF-SEP to visualize neuronal morphology and BDNF release, respectively. Cells were incubated with the transfection mixture for 2.5–3 h at 37°C (95% air and 5% CO₂), rinsed twice with pre-warmed HEPES-buffered solution (HBS; in mM: 135 NaCl, 4 KCl, 1 Na_2_HPO_4_, 2 CaCl_2_, 1 MgCl_2_, 10 glucose, and 20 HEPES; pH 7.35), and returned to fresh Neurobasal medium. Neurons were used for live-cell imaging at 21 DIV.

### Stereotaxic surgery

Adult *Grin1*^flox^ and wild-type mice (P23-35; males and females) were anesthetized with isoflurane (3% vol induction; 1.5% vol maintenance) and secured in a stereotaxic frame (Kopf Instruments, Tujunga, CA). For knockdown of *GRIN1* and expression of BDNF in DG granule neurons, a mixture of AAV5-CamKII-mCherry-cre and cre-dependent AAV-DJ-DIO-BDNF-pHluorin was injected into the DG of the hippocampus (mediolateral, 1.5 mm; anteroposterior, −2.18 mm; dorsoventral, 2.2 mm from bregma) (*92*) at a rate of 0.1 μL/min. A total volume of 1 μL (1:2 ratio) was delivered unilaterally, and the needle remained in place for at least 10 min post-injection. For knockdown of *GRIN1* and expression of BDNF in CA3 pyramidal neurons, a mixture of AAV5-CamKII-mCherry-cre and AAV-hSyn-BDNF-pHluorin was injected into the CA3 (mediolateral, 1.8 mm; anteroposterior, −1.9 mm; dorsoventral, 2.2 mm from bregma) (*92*) at a rate of 50 nL/min. A total volume of 500 nL (1:2 ratio) was injected unilaterally and the needle was left in place for at least 10 min after the injection. Following surgery, mice were monitored for 1 h, returned to their home cages, and allowed to recover for 2–4 weeks before experimentation.

### Acute hippocampal slice preparation

Acute hippocampal slices were prepared as previously described (*22*) with modifications. Two-to-four weeks post-surgery, mice were anesthetized with isoflurane and decapitated. The hippocampus was rapidly dissected, and 300 μm-thick slices were cut using a VT1200S vibratome (Leica Microsystems, Deerfield, IL) in ice-cold NMDG solution containing (in mM): 93 NMDG, 2.5 KCl, 1.25 NaH_2_PO_4_, 30 NaHCO_3_, 20 HEPES, 25 glucose, 5 sodium ascorbate, 2 thiourea, 3 sodium pyruvate, 10 MgCl_2_, and 0.5 CaCl_2_. Slices were transferred to a recovery chamber maintained at 32–34°C and incubated for 30 min in oxygenated artificial cerebrospinal fluid (ACSF) containing (in mM): 124 NaCl, 2.5 KCl, 26 NaHCO_3_, 1 NaH_2_PO_4_, 2.5 CaCl_2_, 1.3 MgSO_4_, and 10 D-glucose. All solutions were continuously bubbled with 95% O_2_ and 5% CO_2_ (pH 7.4). After recovery, slices were maintained at room temperature until imaging experiments.

### Confocal time-lapse imaging

Neuronal cultures were transferred to a temperature- and humidity-controlled imaging chamber and equilibrated in a modified Tyrode’s solution containing (in mM): 119 NaCl, 2.5 KCl, 2 CaCl_2_, 2 MgCl_2_, 25 HEPES, and 30 glucose (pH 7.4). Time-lapse recordings were performed using a 63x/1.4NA Plan-APO oil-immersion objective (Zeiss, Thornwood, NY) on a Zeiss Axio Observer Z1 microscope equipped with a CSU-X spinning-disk confocal scanner (Yokogawa, Wayne, PA). Samples were excited with a laser launch equipped with 50 mW solid-state lasers (488 nm, 561 nm), and fluorescence emission was collected through 525/50 or 629/62 band-pass filters. Images were acquired at 1 min intervals for 60 min using an Orca Flash4 camera (Hamamatsu, Bridgewater, NJ) and Zeiss Zen Blue software. Z-stacks were acquired at a 0.2 to 0.4 μm optical section thickness. BDNF-SEP fluorescence responses were measured in the following order: 3 min baseline recording, 60 min Tyrode’s solution alone or Tyrode’s solution with ketamine (1 μM) or HNK (10 nM), followed by NH_4_Cl alkalization. For pharmacological experiments with NMDA channel blockers, neurons were preincubated with Tyrode’s solution containing AP5 (50 μM) or MK801 (10 μM) for 3 min prior to ketamine application.

### Two-Photon Imaging

Two to four weeks post-surgery, acute hippocampal slices were prepared as described, and slices showing optimal GFP and mCherry expression were selected for imaging. Fluorescence changes were acquired using a Prairie Investigator two-photon microscopy system equipped with a resonant galvo scanning module (Bruker, Billerica, MA). A Chameleon Discovery TPC laser (Coherent, Saxonburg, PA) tuned to 920nm was used for excitation, and signals were collected through GaAsP photomultiplier tubes (Hamamatsu, Bridgewater, NJ) with a 20x/1.0NA objective (Olympus, Tokyo, Japan) at 6.4x zoom. Imaging sites were identified by the presence of axons and large mossy fiber boutons (≥3 μm) in the *stratum lucidum* and captured at 512 x 512 pixel resolution. For presynaptic *GRIN1* knockdown (DG viral injections), mossy fiber terminals co-expressing Cre-dependent BDNF-pHluorin (green) and Cre-dependent mCherry (red) were selected as regions of interest (ROIs). For postsynaptic *GRIN1* knockdown (CA3 viral injections), terminals expressing only non-Cre dependent BDNF-pHluorin were chosen. Boutons expressing Cre-dependent mCherry alone were excluded to avoid retrograde viral spread and unintended *GRIN1* deletion in granule neurons. The time course measurements of BDNF-pH fluorescence were collected under the following sequence: a 1 min ACSF baseline, followed by 5 min of ACSF alone or ACSF containing ketamine (20 μM) or HNK (10 μM). Images were acquired at 1 Hz using T-series software (PrairieView 5.4, Bruker), as modified from (*22, 93*).

For imaging CA3 dendritic spines, acute slices from Thy1-eGFP mice were scanned using the same system, with the addition of a piezoelectric z-axis controller and a 40x/0.8 NA objective (Nikon, Tokyo, Japan). Imaging sites were identified by dendritic morphology in the *stratum lucidum* or *stratum radiatum*. Spines, primarily located on second-order apical dendrites, were selected as ROIs from 512 × 512-pixel images (78 × 78 μm field of view; pixel size: 0.152 μm; *z*-step: 0.5 μm). Two Z-stacks were collected for each slice: one immediately after initiating ACSF perfusion (baseline) and after 30 min of continuous perfusion with ACSF alone, ACSF + ketamine (20 μM), or ACSF + HNK (10 μM).

### Image and statistical analysis

#### Electron microscopy

All analyses were done by persons who were blind to experimental groups. Ultrathin sections from 3 mice in each experimental condition was analyzed at the tissue-plastic interface to minimize the difference in antibody penetration (*40, 94*). Immunolabeled profiles were classified using defined morphological criteria. Dendritic profiles contain cytoplasm with mitochondria, microtubules, and agranular reticulum and were postsynaptic to axon terminal profiles identified by the presence of vesicles. Mossy fiber pathway terminals are identified by their large size (1–2 μm in diameter) and often contact multiple dendritic spines of the CA3 pyramidal cell dendrites (*95*). For quantification, 50 GluN3A-labeled mossy fiber terminals from randomly selected 3,555 nm^2^ fields in CA3 were analyzed, and GluN3A-immunogold particles were manually counted within mossy terminals and their postsynaptic spines.

#### BDNF-SEP

Asynchronous BDNF release from neuronal cultures is well documented and was quantified as the number of events, rather than changes in fluorescence intensity (*16, 91*). All images were background-subtracted, corrected for XY drift using the StackReg plugin in ImageJ, and sequential frames were aligned to ensure accurate tracking of BDNF release events and to prevent motion artifacts from influencing fluorescence measurements. ROIs were selected based on mCherry-labeled (red) boutons to restrict analysis to BDNF release events originating from mossy fiber terminals. Detection of fluorescence changes indicating BDNF release within each ROI was performed using a custom-modified graphical user interface (MATLAB; FluoroSNNAP; University of Pennsylvania), which automatically determined fluorescence thresholds, filtered particle size, and excluded events outside the defined ROI. Fluorescence intensity for each detected event was calculated by measuring the change from baseline fluorescence using a custom Time Series Analyzer plugin in ImageJ. Each event underwent signal-to-noise ratio (SNR) analysis, and ΔF was defined as the difference in peak fluorescence (F_peak_) and baseline fluorescence (F_baseline_). Release events counted only if mean ΔF was >5 SD above the baseline mean. Baseline fluorescence (F_baseline_) was calculated from three frames acquired over a 3-min period prior to treatment, and ΔF was calculated as the difference between the peak fluorescence (F_peak_) and F_baseline_. Because each FOV contained a different number of boutons, event frequency was normalized by dividing the total number of BDNF release events by the number of boutons within that FOV. Each imaging session ended with alkalization using 50 mM NH_4_Cl to confirm localization of BDNF-pH and to verify cell responsiveness. BDNF trafficking particles were excluded from the analysis, and only NH_4_Cl-responsive cells were included.

#### BDNF-pHluorin

All images were background-subtracted, corrected for XY drift during acquisition using the Movie Aligner plugin in ImageJ (NIH, Bethesda, MD), and aligned frame-by-frame previously as described (*96*) to ensure accurate tracking of BDNF release events and prevent motion artifacts from influencing fluorescence measurements. ROIs were identified manually, and fluorescence intensity was quantified for each ROI relative to its baseline. The difference between F_peak_ and F_baseline_ was used to calculate ΔF, and values were reported as ΔF/F of the BDNF-pHluorin signal (*22*) using ImageJ (Time series analyzer, NIH).

#### Quantitative spine analysis

Dendritic branches from acquired images were processed using Huygens deconvolution software (Scientific Volume Imaging, Hilversum, Netherlands) to enhance SNR ratio and spatial resolution. Following deconvolution, spine density was quantified using Restoration Enhanced Spine and Neuron Analysis (RESPAN) by selecting dendritic segments or ROIs of fixed length and width for each neuron, with each ROI containing at least 45 spines (*97, 98*). Dendritic arbors were segmented to identify spine heads and necks from eGFP fluorescence using an adaptive, self-configuring nnU-Net architecture (*99*). Subsequent processing in RESPAN, using the Model 3 pre-trained segmentation model available through the official GitHub repository, generated three-dimensional spine detection outputs that were used to quantify spine density, reported as spines per micrometer of dendrite. To minimize false positives, spines were retained only if their volumes ranged from 0.03 to 15 µm^3^ and their distances from the parent dendrite were below the defined threshold. Dendrite segments with total volumes < 3 µm^3^ were excluded. Individual spine numbers were measured at baseline, 0 min, and after 30 min. Across experimental groups, 375–925 spines were analyzed per condition. Automated spine detection and density measurements were independently validated by manual spine counts conducted on both original and deconvolved images.

### Data analysis

All analyses were performed with the experimenter blinded to experimental conditions. Statistical comparisons were conducted using unpaired two-tailed Student *t-*tests or ordinary one-way ANOVA, followed by Tukey *post-hoc* tests, as appropriate, using GraphPad Prism v10 (Graphpad Software, Inc., San Diego, CA). A significance threshold of *P* < 0.05 was applied. Data are presented as mean ± SEM unless otherwise indicated. Asterisks denote values significantly different from control (CTL): \**P* < 0.05, \*\**P* < 0.01, \*\*\**P* < 0.001, \*\*\*\**P* <0.0001.

## Acknowledgments

We thank Dr. Hyungju Park (Korea Brain Research Institute, Daegu, Korea) for generously providing plasmids for the BDNF-pH viral vector. We thank Mark Koenis (Huygens; Scientific Volume Imaging, Hilversum, Netherlands) and Luke Hammond (RESPAN) for their help with data processing for spine quantification. We also thank Ms. June Chan of the Neuroanatomy EM core, Feil Family Brain and Mind Research Institute, for technical assistance.

## Author contributions

JP, ILAR, and HCH conceived and designed the study. Material preparation, data collection, and analysis for Figures 1, 2E-F, and 3–5 were performed by ILAR. KP, HA, and SZ contributed to data collection and analysis for Figure 2A–D. KFH contributed to data analysis for Figure 5 and assisted with figure formatting across all figures. Custom macros were written by KWJ. Figures were prepared by ILAR, KFH, and JP. The original draft of the manuscript was written by JP, with additional review and editing by JP, HCH, and TAM. All authors approved the final manuscript.

## Supplementary Materials

**Supplementary Video 1 (attached)**: Ketamine (1 μM) evokes rapid release of BDNF Representative time-lapse video of rat hippocampal neurons (21 DIV) expressing BDNF-SEP and mCherry (not shown), perfused with ketamine (1μM), showing discrete BDNF exocytotic events (green puncta) from mossy fiber terminals over a 60-min imaging period.

### Supplementary Figures

**Fig. S1:**
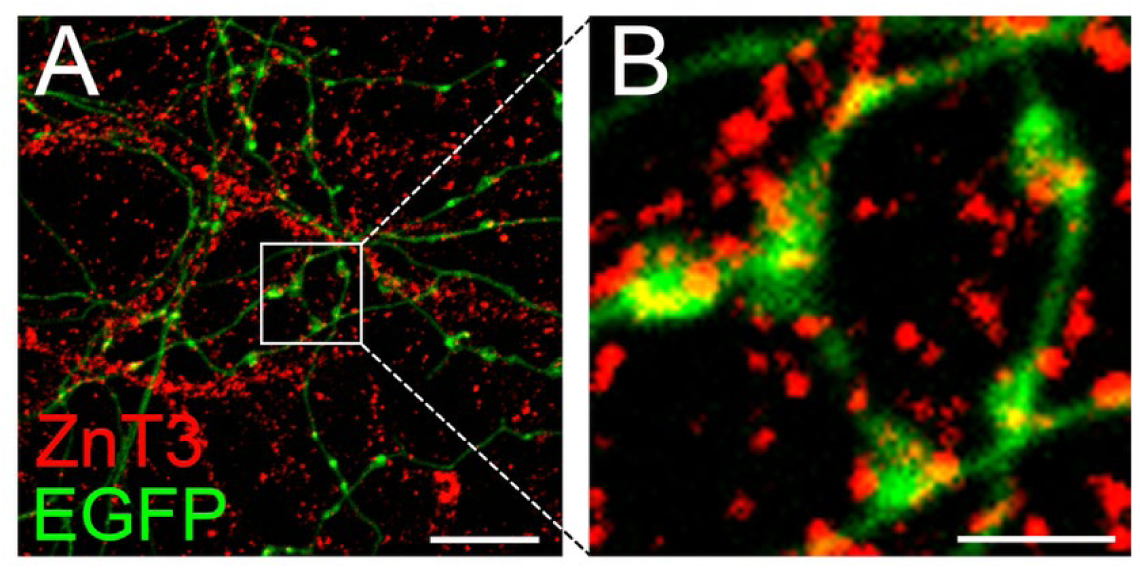
Zinc transporter 3 (ZnT3) is expressed in mossy fiber terminals. Immunofluorescence images showing axons (**A**) with close-up of mossy fiber terminals (**B**) labeled with ZnT3. Scale bars = 10 μm (**A**) and 5 μm (**B**).

**Fig. S2.**
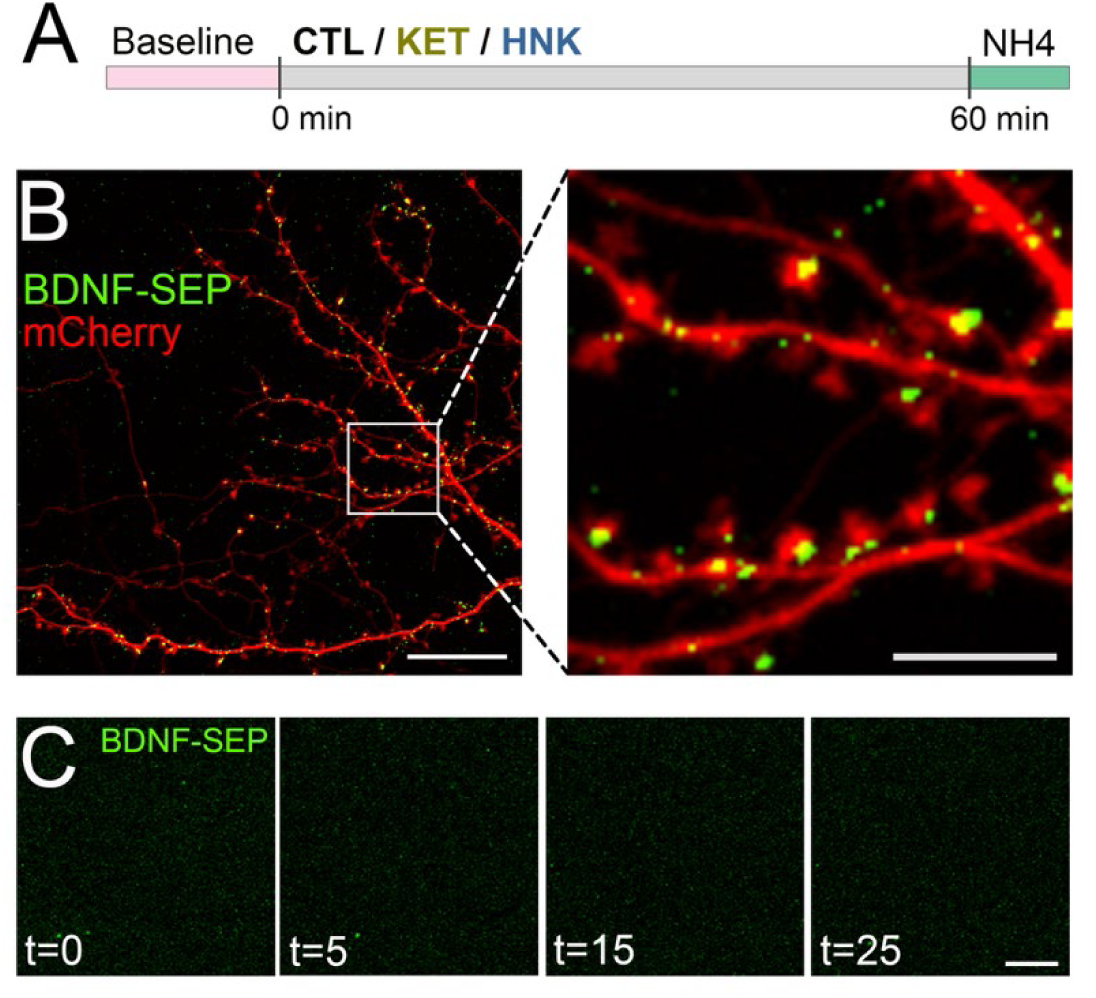
Ketamine or HNK does not release BDNF from hippocampal dendritic spines. Rat hippocampal neuron cultures (21 DIV) transfected with BDNF-SEP and mCherry were perfused with 1μM ketamine, 10nM HNK, or control buffer and imaged as indicated in (**A**). Representative images showing dendritic arbor of a control neuron (**B,** *left*) with close-up of dendritic spines (**B**, *right*) after alkalinization with BDNF-SEP pools in spines. Representative images showing single BDNF exocytotic events before (**C**, t = 0) and after (**B**, t = 5–25) ketamine application, with no release of BDNF detected. Scale bar = 10 μm (**B,** *left*; **C**) and 5 μm (**B,** *right*).

